# High Estradiol reduces adult neurogenesis but strengthens functional connectivity within the hippocampus during spatial pattern separation in adult female rats

**DOI:** 10.1101/2022.12.22.521710

**Authors:** Shunya Yagi, Stephanie E. Lieblich, Liisa A.M. Galea

## Abstract

Adult neurogenesis in the dentate gyrus plays an important role for pattern separation, the process of separating similar inputs and forming distinct neural representations. Estradiol modulates neurogenesis and hippocampus function, but to date no examination of estradiol’s effects on pattern separation have been conducted. Here, we examined estrogenic regulation of adult neurogenesis and functional connectivity in the hippocampus after the spatial pattern separation task in female rats. Ovariectomized Sprague-Dawley rats received daily injections of vehicle, 0.32µg (Low) or 5 µg (High) of estradiol benzoate until the end of experiment. A single bromodeoxyuridine (BrdU) was injected one day after initiation of hormone or vehicle treatment and rats were tested in the delayed nonmatching to position spatial pattern separation task in the 8-arm radial maze for 14 days beginning two weeks after BrdU injection. Rats were perfused 90 minutes after the final trial and brain sections were immunohistochemically stained for BrdU/neuronal nuclei (NeuN) (new neurons), Ki67 (cell proliferation), and the immediate early gene, zif268 (activation). Results showed that only high estradiol reduced the density of BrdU/NeuN-ir cells and had significant inter-regional correlations of zif268-ir cell density in the hippocampus following pattern separation. Estradiol treatment did not influence pattern separation performance or strategy use. These results show that higher doses of estradiol can reduce neurogenesis but at the same time increases correlations of activity of neurons within the hippocampus during spatial pattern separation.

## 1. Introduction

New neurons are continuously generated in the subgranular zone of the dentate gyrus (DG) in the hippocampus in adulthood. These new neurons play a critical role in pattern separation (Clelland et al., 2009), a process that enables discrimination and episodic memory by separating similar memory patterns to make distinct neural representations (Marr, 1971). Estradiol modulates both neurogenesis and spatial ability in a dose-dependent manner (reviewed in Duarte-Guterman et al., 2015) but to our knowledge, no studies have examined whether estradiol can modulate pattern separation. Therefore, the present study aimed to elucidate the effects of two different doses of estradiol on pattern separation, adult neurogenesis and neural activation in the hippocampus. We hypothesized that estradiol would modulate the ability for pattern separation, adult neurogenesis and zif268 activation in the hippocampus.

## 2. Methods

### 2.1. Subjects

Twenty-five two-month-old female Sprague Dawley rats were purchased from Charles River Canada (St-Constant, Quebec, Canada). Rats were initially pair-housed for two weeks after arrival and single-housed afterward throughout entire experiment. All experiments were carried out in accordance with Canadian Council for Animal Care guidelines and were approved by the animal care committee at the University of British Columbia (A20-0147). All efforts were made to reduce the number of animals used and their suffering during all procedures.

### 2.2. Experimental design

All animals were handled every day for 2 minutes beginning one week after arrival. All rats were bilaterally ovariectomized two weeks after arrival (Fig. 1A). Ovariectomized rats received daily subcutaneous injections of 0.32 µg (Low) or 5 µg (High) estradiol benzoate, or vehicle in 0.1ml of sesame oil beginning one week after surgery (Day 0) until the end of experiment. One day after the initiation of hormone or vehicle treatment, one intraperitoneal injection of bromodeoxyuridine (BrdU; 200mg/kg; Sigma-Aldrich, Oakville, ON, Canada) was administered to all animals two hours after the second hormone/vehicle treatment (Day1). Four days after BrdU injection, all animals were food restricted and maintained their weight at 87-92% of their original weight throughout entire behavioral testing. On Day 8 to Day 10, rats were habituated to the RAM and on Days 11-13, rats were shaped (food reward placed on arms of radial arm maze) for 5 minutes each day. Following shaping, all rats were tested in the delayed nonmatching to position (DNMP) RAM task for 14 days (Day14-27; Fig. 1B) one hour after hormone/vehicle treatment, which was followed by an activation and probe trial (Day28; Fig. 1C). On the last day (Fig. 1D), rats received one activation trial (same procedure as the spatial pattern separation task). Rats then received a probe trial to determine whether they were idiothetic or place strategy users 80 minutes after the activation trial. Ninety minutes after the activation trial, all rats were perfused and underwent tissue collection. Brain sections were immunohistochemically stained for Ki67, zif268 and BrdU/NeuN (all methods are described in detail in the Extended Experimental Procedures). The present study analyzed the dorsal and ventral hippocampus separately as the dorsal hippocampus is important for spatial learning and memory, whereas the ventral hippocampus is important for regulation of anxiety and stress response (Kjelstrup et al., 2002; Moser et al., 1993).

**Fig. 1.**
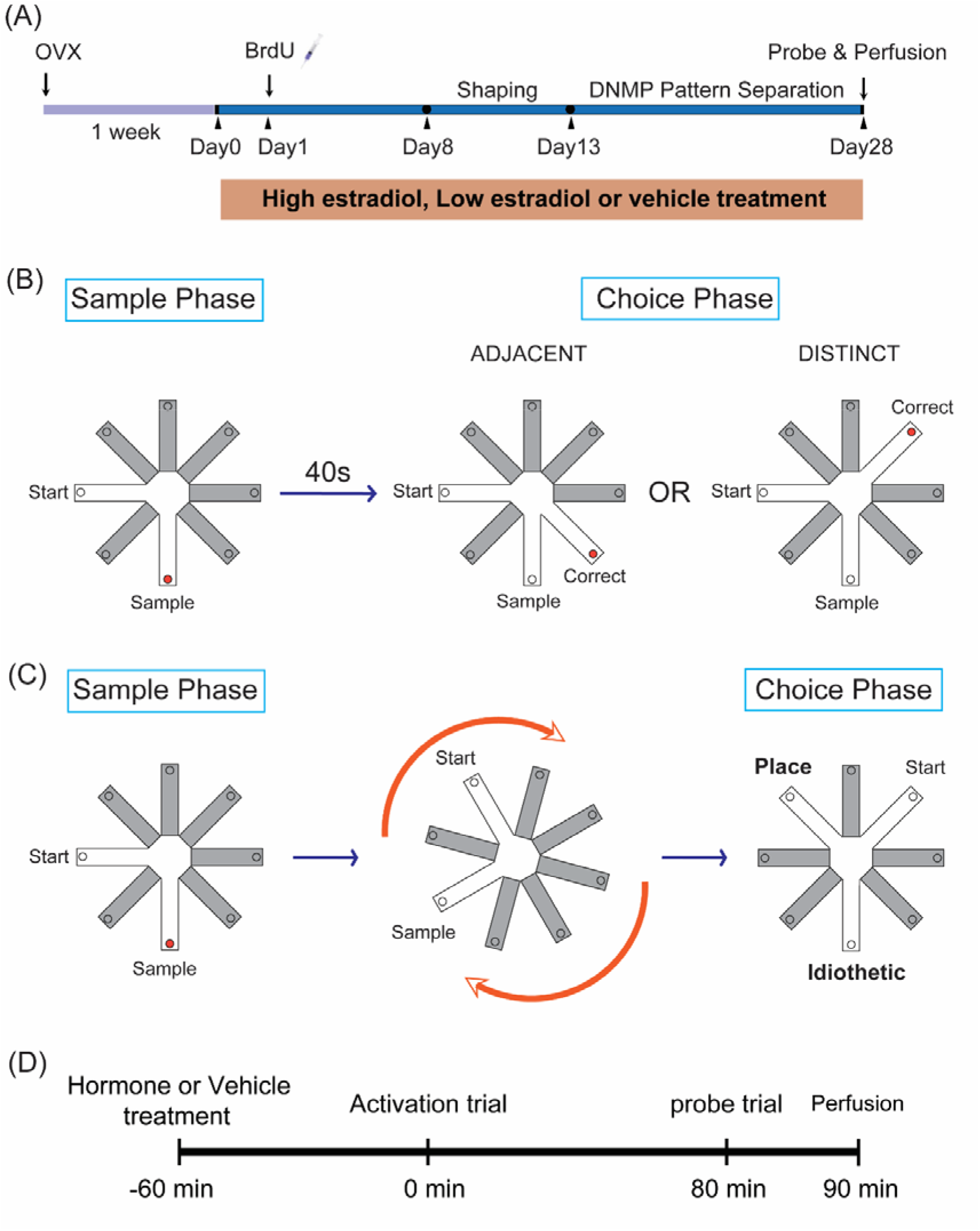
Experimental design of the spatial pattern separation task in ovariectomized rats. (A) Timeline of the experiment. (B) The spatial pattern separation task utilized the delayed non-match to position task in the radial arm maze, and rats received four trials each day (two trials of each separation arm pattern – distinct and adjacent). Each trial consists of sample phase and choice phase where rats were tested in their ability to discriminate the newly-opened arm during choice phases. During the sample phase, a rat was placed on the start arm and allowed to retrieve a food reward (indicated with orange circle in Fig. 1B) at the end of Sample arm. During the choice phase, the additional/correct arm, but not the sample arm, was baited (Correct arm). (C) Rats were classified as a place strategy user or an idiothetic strategy user based on their arm choice during the choice phase of probe trial. After the sample phase of the probe trial, the maze was rotated and the start arm was moved to a new location and two choice arms were presented. One choice arm (Idiothetic arm) was held the same position as the sample arm relative to the extra-maze cues and the other choice arm (Place arm) was perpendicularly right from the new start arm. A place strategy user will choose the newly opened arm by using extra-maze cues, whereas an idiothetic strategy user will choose the sample arm with ignoring extra-maze cues as the orientation of start-Place arm pair (90°) during the choice phase was the same as that of start-sample arm pair (90°) during the sample phase. (D) Timeline on the last day. All rats were perfused 90 minutes after the activation trial.

### 2.3. Statistical analysis

All analyses were conducted using Statistica (Statsoft Tulsa, OK, USA) and significance level was set at α = 0.05. The percentage of correct choices during ADJACENT or DISTINCT trials were each analyzed using two-way analysis of variance (ANOVA), with strategy choice (spatial, idiothetic) and treatment (Vehicle, Low dose, High dose) as between-subject variables. Chi-square analyses were used for strategy choice across the treatment. Two-way ANOVAs were used for each dorsal (d) and ventral (v) DG to analyze the density of Ki67-ir cells and BrdU/NeuN-ir cells, and for each subregion of the hippocampus (dDG, vDG, dCA3, vCA3, dCA1, vCA1) to analyze the density of zif268-ir cells with strategy choice and treatment as between-subject variables. Post-hoc tests utilized the Neuman-Keuls procedure. As a measure of functional connectivity, Pearson product-moment correlations were also calculated between the density of zif268-ir cells among the hippocampus subregions and a brain network map for each treatment was calculated with coefficient of correlations (r) for each inter-regional correlation. The coefficient of variation (r^2^) for each inter-regional correlation for each treatment group was calculated to measure the proportion of variance accounted for between the two subregions of the hippocampus, which was used as a measure of the strength of connectivity (Liu et al., 2022). The r^2^ values were analyzed using one-way ANOVA with treatment as the between-subject variables. In addition, Pearson product-moment correlations were calculated between the density of BrdU/NeuN-ir cells in the dDG or vDG and the density of zif268-ir cells in each subregion of the hippocampus. A priori comparisons and Pearson product-moment correlations were subjected to Bonferroni corrections.

After the sample phase of the probe trial, the maze was rotated and the start arm was moved to a new location and two choice arms were presented. One choice arm (Idiothetic arm) was held the same position as the sample arm relative to the extra-maze cues and the other choice arm (Place arm) was perpendicularly right from the new start arm. A place strategy user will choose the newly opened arm by using extra-maze cues, whereas an idiothetic strategy user will choose the sample arm with ignoring extra-maze cues as the orientation of start-Place arm pair (90°) during the choice phase was the same as that of start-sample arm pair (90°) during the sample phase. (D) Timeline on the last day. All rats were perfused 90 minutes after the activation trial.

## 3. Results

### 3.1. There was no significant effect of estradiol on strategy use or on the ability for separating either adjacent or distinct patterns

Neither chronic administration of High nor Low estradiol affected strategy use nor the percentage of correct arms chosen regardless of the pattern (adjacent or distinct) compared to vehicle treatment (p’s > 0.345; Supplemental Fig. 1A-B).

### 3.2. High estradiol reduced the density of BrdU/NeuN-ir cells in the dDG

High estradiol-treated rats had significantly less density of BrdU/NeuN double-ir cells in the dDG compared to vehicle-treated rats (p = 0.011, Cohen’s d = 1.846) and a trend compared to Low estradiol-treated rats (p = 0.065, Cohen’s d = 1.045) [main effect of treatment: F(2, 17) = 4.621, p = 0.025, partial η^2^ = 0.352; Fig. 2A]. There were no other significant main or interaction effects on the density of BrdU/NeuN-ir cells (p > 0.299) or Ki67-ir cells (p’s > 0.116; Supplemental Fig. 1C-D) in the DG.

**Fig. 2.**
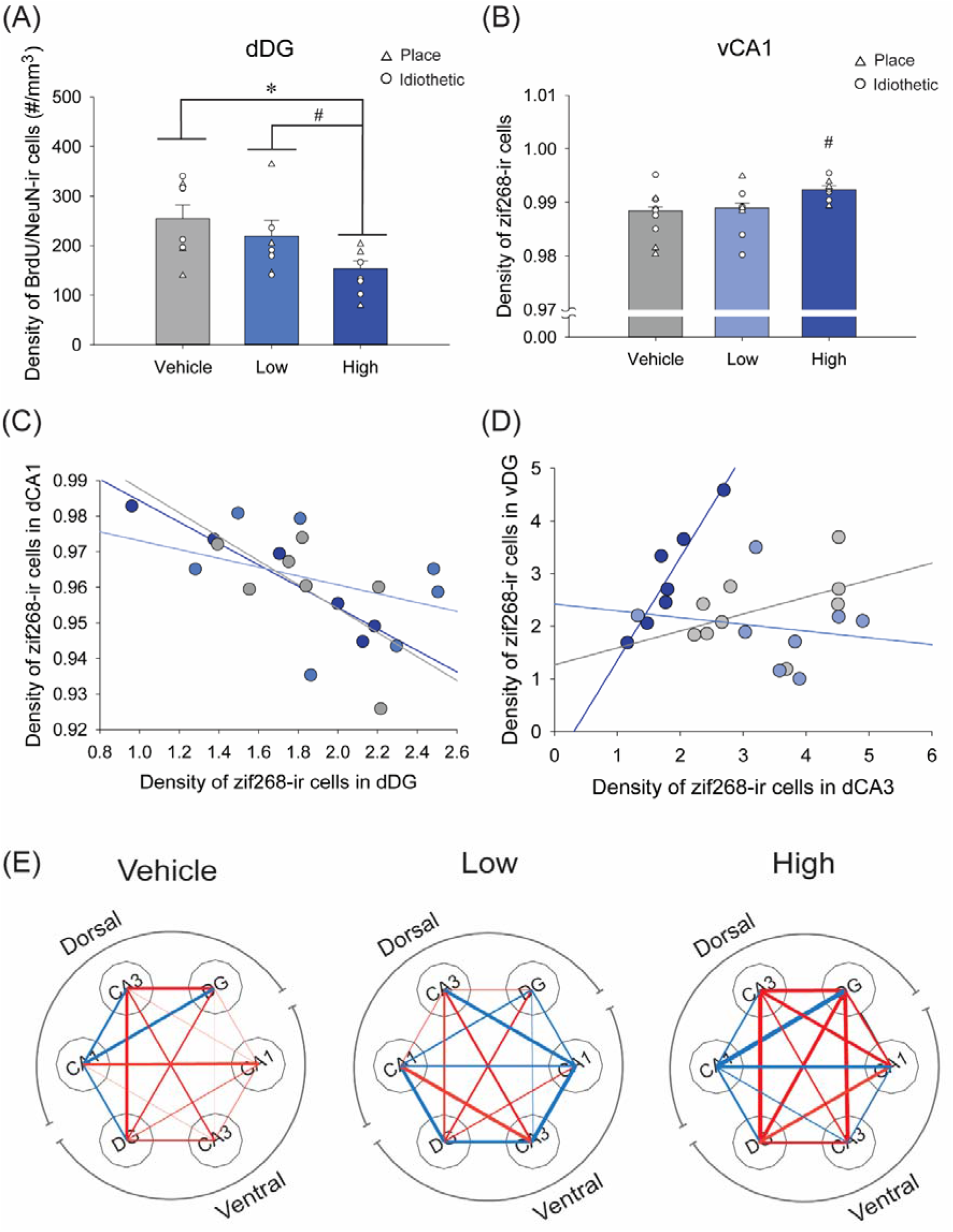
(A) Mean density of bromodeoxyuridine (BrdU)/neuronal nuclei (NeuN)-immunoreactive (ir) cells in the dorsal (d) dentate gyrus (DG). High estradiol-treated rats had a significantly less density of BrdU/NeuN-ir cells and Low estradiol-treated rats had a trend of less density of BrdU/NeuN-ir cells in the dDG compared to vehicle-treated rats. (B) Mean density of zif268-ir cells in the ventral (v) Cornu Ammonis (CA) 1. High estradiol-treated rats had a trend of greater density of zif268-ir cells in the vCA1 compared to Low estradiol-treated rats and compared to vehicle-treated rats. (C-D) Inter-regional correlations of the density of zif268-ir cells in vehicle-treated (grey), Low estradiol-treated (light blue) and High estradiol-treated rats (dark blue). High estradiol-treated rats had a negative correlation between the dCA1 and the dDG (C) and a positive correlation between the vDG and the dCA3 (D). (E) Brain network maps were generated with correlations with red lines indicating positive correlations and blue lines indicating negative correlations with wider lines indicating larger coefficients in vehicle-treated (Vehicle), Low estradiol-treated (Low) and High estradiol-treated (High) rats. High estradiol treated rats had greater overall r^2^ values compared to other groups. Error bars represent ± SEM. * indicates p < 0.05 and # indicates 0.05< p < 0.10.

### 3.3. zif268 expression in the vCA1 was greater in High estradiol-treated rats than the other groups

High estradiol-treated rats tended to have a greater density of zif268-ir cells in the vCA1 compared to vehicle-treated rats and Low estradiol-treated rats [vehicle: p = 0.058, Cohen’s d = 1.320; Low: p = 0.070, Cohen’s d = 1.158; main effect of treatment: F(2, 18) = 3.167, p = 0.066, partial η^2^ = 0.352; see Fig. 2B]. There were no main or interaction effects on the density of zif268-ir cells in the hippocampus (p’s > 0.222).

### 3.4. High estradiol-treated rats had distinct inter-regional correlations compared to the other groups

For both estradiol-treated groups there were significant intra-hippocampal correlations of zif268 activity between the six regions, but there were no significant correlations in the vehicle group (3 in High, 1 in Low, 0 in Vehicle). Intriguingly, all the three correlations in the High estradiol-treated group involved the DG (2 involved dDG and 1 involved vDG) but not in the Low estradiol-treated group. However, after applying Bonferroni corrections, the High estradiol-treated rats had two significant correlations in the density of zif268-ir cells between the dDG and the dCA1 [r(7) = -0.963, p = 0.002; Fig. 2C] and the vDG and the dCA3 [r(6) = 0.940, p = 0.002; Fig. 2D]. There were no significant inter-regional correlations in the hippocampus in the other two groups after Bonferroni corrections (p’s > 0.030). Furthermore, High estradiol-treated rats had greater coefficients of variation (mean r^2^ = 0.445 ± 0.073) in the hippocampus compared to Low estradiol-treated rats (mean r^2^ = 0.240 ± 0.054) and vehicle-treated rats (mean r^2^ = 0.153 ± 0.043) [Low estradiol: p = 0.016, Cohen’s d = 0.827, Cohen’s d = 1.265; vehicle: p = 0.003; main effect of treatment: F(2, 42) = 6.746, p = 0.003, partial η^2^ = 0.243]. There was no significant difference between Low estradiol-treated rats and vehicle-treated rats (p = 0.292).

In addition, there were no significant correlations between the density of BrdU/NeuN-ir cells and the density of zif268-ir cells in any hippocampal subregions after Bonferroni correction (p’s > 0.035).

## 4. Discussion

### 4.1. High levels of estradiol decreased adult-born neurons in the dDG

The present study demonstrates that high levels of estradiol decreased the density of new neurons (BrdU/NeuN-ir) in the dDG compared to vehicle-treated rats, consistent with past studies in female rats that did not undergo spatial testing (Chan et al., 2014). Estradiol can upregulate or downregulate neurogenesis dependent on the timing of estradiol and BrdU injection (B. K. Ormerod et al., 2003b) and to cognitive training (McClure et al., 2013). Because in the present study we injected BrdU 26 h after the first injection of high estradiol benzoate, this likely lead to the reduction in survival of new neurons, due to lower cell proliferation at that time (B. K. Ormerod et al., 2003b). The ability of estradiol to reduce cell proliferation after 24-48 h in females is due to estradiol’s effects to increase adrenal steroids, but not on estradiol’s effects on NMDA receptor activation (B. K. Ormerod et al., 2003a). Further research is warranted to elucidate molecular, cellular and systemic mechanisms underlying the effect of timing of estradiol on adult neurogenesis in females and how cognitive training may influence this.

### 4.2. Estradiol altered patterns of neural activation and functional connectivity in the hippocampus

Previous studies demonstrate that estradiol increases CA1 dendritic spine density through activation NMDA receptors and increases synaptic excitability through activation of AMPA receptors (Wong and Moss, 1992; Woolley and McEwen, 1994). Thus, it is possible that greater neural activation in the vCA1 in the present study is due to the long-term effect (increasing dendritic spines) or the short-term effect (increasing excitability) of estradiol. However, in this study we used 29 days of treatment with estradiol, while others have used acute treatment of estradiol on the electrophysiological properties of CA1 neurons, thus the influence of chronic estradiol treatment on electrophysiological properties are still unknown. The present study also demonstrates that high estradiol rats have an inverse association of the neural activation between the dDG and dCA1, and neural coactivation between vDG and dCA3. Furthermore, high estradiol rats had greater overall amount of variance accounted for across inter-regional correlations in the hippocampus compared to the other groups. These findings are important as it suggests that chronic high levels of estradiol enhance functional connectivity among hippocampal subregions during spatial learning, although this did not affect learning outcomes. Previous work in humans and rodents demonstrate that functional connectivity in the hippocampus decreases with age, which is associated with spatial learning and memory (Dalton et al., 2019; Liang et al., 2020). Taken together, it is possible that high estradiol treatment can restore age-related decline in functional connectivity in the hippocampus, and potentially improve hippocampus-dependent cognition in aged subjects.

### 4.3. Strategy use was not altered by estradiol dose

Estradiol dose did not affect strategy use in the present study and this is inconsistent with findings from other studies examining strategy use across the estrous cycle (Korol et al., 2004). The present study also failed to demonstrate significant strategy differences in zif268 activation in the dorsal hippocampus or significant correlations between the density of zif268-ir cells and the density of new neurons in the dorsal hippocampus. These results indicate that the hippocampus was equally recruited during the activation trial between the two strategy users. The present study used daily injections of 5μg or 0.32μg of estradiol benzoate into young ovariectomized rats, which gives serum concentrations of estradiol equivalent to proestrous or diestrous females, respectively (Becker and Rudick, 1999). However, repeated exogenous estrogens may influence adult neurogenesis and cognition differently than naturally fluctuating estrogens. Proestrus is associated with a hippocampus-dependent place strategy, while diestrus is associated with striatum-dependent response strategy (reviewed in Yagi and Galea, 2019). Furthermore, low estradiol phase females perform better at spatial learning tasks in the Morris water maze compared to high estradiol phase females (Galea et al., 1995; Warren and Juraska, 1997) and thus perhaps other ovarian hormones work to influence strategy use.

In the present study we also saw no significant effects of chronic estradiol benzoate treatment on spatial pattern separation performance, inconsistent with past work that indicated that low and high chronic estradiol benzoate modulated working memory in the spatial radial arm maze (Holmes et al., 2002). Consistent with other studies, the present study shows that long-term exposure to exogenous estradiol loses the enhancing effect of estradiol on cell proliferation(reviewed in Duarte-Guterman et al., 2015). Therefore, further research is needed to elucidate complex effects of estradiol on cognition and underlying mechanisms.

## Conclusion

The present study demonstrates that estradiol benzoate modulates neurogenesis and activity within the hippocampus of young adult female rats in a dose dependent manner, with no significant effect of estradiol on pattern separation performance. These findings highlight the importance of considering the hormonal status when studying brain connectome and hippocampal neural plasticity.

## Availability of data and materials

The datasets used and/or analyzed during the current study are available from the corresponding author on reasonable request.

## Authors’ contribution

**SY:** Conceptualization, Methodology, Data curation, Writing-Original draft preparation, Visualization. **SL:** Data collection. **LG**: Conceptualization, Methodology, Analysis, Writing-Reviewing and Editing, Supervision.

## Acknowledgment

We would like to thank Carmen Chow for the exceptional technical assistance with this work.

## Extended Experimental Procedures

### 1. Animal husbandry

Rats were housed in opaque polyurethane bins (48 × 27 × 20 cm) with paper towels, polyvinylchloride tube, and cedar bedding. Rats have free access to water and normal lab chow, and maintained under a 12 : 12 hour light/ dark cycle (7A.M. light-on).

### 2. Apparatus

The radial arm maze (RAM) had an octagonal center platform (36 cm in diameter) and 8 arms (53 cm long × 10 cm wide) that were set 80 cm above the floor in the center of a dimly lit room. Large extramaze cues were placed on all four walls of the room and were not moved throughout the study. Metal gates were used to block entries to arms.

### 3. Ovariectomy and hormone replacement

Rats were anesthetized using an initial flow rate of 5% isofluorane (Boxter Corp., Mississauga, ON, Canada) and 2-3% during surgery. 35 mg/kg Ketamine (Merial Canada Inc, Baie-d’Urfe, QC, Canada) and 1.5 mg/kg Xylazine were administered intraperitoneal, and 4mg/kg Marcaine HCL and 5mg/kg Anafen were administered subcutaneously (s.c.) before surgery. To prevent dehydration, 5 ml Lactated Ringer Solution (Braun Medical Inc, Scarborough, ON, Canada) was injected s.c. After a recovery phase of six days, the rats were divided into three groups, eight rats for high dose estradiol group, eight rats for low dose estradiol group and nine rats for vehicle treated group. High dose estradiol group received a s.c. injection of 5 μg estradiol benzoate (E2B: Sigma) in 0.1 ml sesame oil, low dose estradiol group received 0.32 μg E2B in 0.1ml sesame oil and vehicle treated group received 0.1 ml sesame oil for 29 consecutive days including the probe trial.

### 4. Habituation, shaping and behavioral testing for spatial pattern separation

During habituation, rats were placed on the center platform and allow them to freely explore all the arms for 10 minutes. During the first day of shaping, three quarters of Froot Loops (Kellog’s) were located along each arm at equidistant intervals and a quarter was placed in a cup (3 cm in diameter) at the end of arms. During the second and the third day of shaping, each arm was baited only with a quarter of Froot Loops placed in each cup.

Rats received four trials of delayed non-match place (DNMP) version of pattern separation task in the radial arm maze (RAM) each day (two trials of each separation pattern) for 14 consecutive days (56 trials total with 28 trials of each separation). The first trial of each day began between 10 a.m. and 11 a.m two hours after hormone or vehicle treatment.

Every rat received one trial before the first rat began their next trial to maximize the intervals between each trial. One trial consisted of two phases, a sample phase and a choice phase (40 seconds interval between the two phases). Rats were tested in their ability to discriminate the newly-opened arm during choice phases. During the sample phase, a start arm and a sample arm were open and all the other arms were closed. A rat was placed on the start arm and the rat was allowed to visit the sample arm and retrieve a quarter of Froot Loop (reward). Rats were returned to their cage from the maze after spending ten seconds in the sample arm after eating the reward or exiting the sample arm. During the choice phase, all arms were closed except the start, sample and an additional arm (correct arm). The additional/correct arm, but not the sample arm, was baited during the choice phase. Rats that made incorrect choices (entry to the sample arm or start arm) were permitted to self-correct and retrieve the reward from the correct arm. Rats were retrieved from the maze after rewarded or after one minute had passed and returned to the colony room. During the 40-second interval between a sample phase and a choice phase, the maze was rotated to minimize the ability of rats to reach the correct arm by utilizing intra-maze cues such as odor. After the rotation, the location of the start and sample arms relative to extra-maze cues, but not the arms themselves, were held constant during each trial. The RAM was wiped with distilled water after each rat.

Two sets of sample-correct arm pairs were used in this study, ADJACENT and DISTINCT. Correct arms during ADJACENT trials were 45° away from the sample arm and correct arms during DISTINCT trials were 135° away from the sample arm. Start arms were located perpendicular to either the correct or sample arms (Figure. 1B). Sample-correct-start arm combinations were pseudo-randomly chosen for each day from the pool of possible combinations so that overlaps in the presentation of arms were minimized both within each day and across the entire experiment. The order of rats to be tested each day was randomized every day throughout this experiment.

### 5. Probe trial

On Day 28, rats received hormone or vehicle injection in the morning between 9 a.m. and 10 a.m. One hour after hormone or vehicle treatment, rats received one set of activation trial (same procedure as DNMP version of spatial pattern separation task described previously). Following 80 minutes after the activation trial, rats received a probe trial to determine whether they were response strategy users (relying more on idiothetic response cues) or place strategy users (relying more on spatial extra-maze cues). A probe trial consisted of a sample phase and a choice phase with 40 seconds of interval between the two phases. A sample phase of the probe trial has the same rules as a sample phase of testing trials. The start arm during sample phase was perpendicular to the sample arm that was located to the right from the start arm. After the sample phase, all arms were blocked off and the maze was rotated. The start arm was moved to a new position and two choice arms (135° away from each other) were opened after the rotation. One choice arm (Idiothetic arm) was held the same position as the sample arm relative to the extra-maze cues and the other choice arm (Place arm) was perpendicularly right from the new start arm. A place strategy user chose the newly opened arm by using extra-maze cues, while an idiothetic strategy user chose the sample arm with ignoring extra-maze cues as the orientation of start-Place arm pair (90°) during the choice phase was the same as that of start-sample arm pair (90°) during the sample phase (Fig. 1C). Immediately after the probe trial, rats were perfused (about 90 minutes after the activation trial).

### 6. Perfusion and tissue processing

Rats were administered an overdose of sodium pentobarbitol (500 mg/kg, i.p.) and perfused transcardially with 60 ml of 0.9% saline followed by 120 ml of 4% formaldehyde (Sigma-Aldrich). Brains were extracted and post-fixed in 4% formaldehyde overnight, then transferred to 30% sucrose (Fisher Scientific) solution for cryoprotection and remained in the solution until sectioning. Brains were sliced into 30 μm coronal sections using a Leica SM2000R microtome (Richmond Hill, Ontario, Canada). Sections were collected in series of ten throughout the entire rostral-caudal extent of the hippocampus and stored in anti-freeze solution consisting of ethylene glycol, glycerol and 0.1M PBS at -20°C.

### 7. Immunohistochemistry

#### 7.1. Ki-67 DAB staining

Brain sections were rinsed overnight with 0.1 MPBS at 4 °C. The tissue was incubated in 0.6% H_2_O_2_ for 30 minutes and then incubated in primary antibody solution containing 1:1000 rabbit anti-Ki67 antibody (Vector Laboratories), 0.3% Triton-X, and 3% normal goat serum (NGS; Vector Laboratories) in 0.1 M PBS for 24 hours at 4 °C. Following rinsing the sections four times, the sections were incubated in secondary antibody solution consisting of 1:1000 goat anti-rabbit biotinylated IgG (Vector Laboratories, Burlington, ON, Canada) in 0.1 M PBS for 24 hours at 4 °C. The sections were then incubated in AB solution (Vector Laboratories) for 1 hour at room temperature. The sections were then visualized with diaminobenzidine (DAB; Sigma) solution and mounted onto microscope slides, followed by dehydrated, cleared with xylene and cover-slipped with Permount (Fisher Scientific; Ottawa, ON, Canada).

#### 7.2. Zif268 DAB staining

Brain tissue was rinsed overnight with 0.1 MPBS at 4 °C. The tissue was incubated in 0.6% H_2_O_2_ for 30 minutes and then incubated in primary antibody solution containing 1:1000 Rabbit anti-Erg-1 (Milli-pore; MA, USA) or anti-cFos (Milli-pore; MA, USA), 0.04% Triton-X, and 3% normal goat serum (NGS; Vector Laboratories) in 0.1 M PBS for 24 hours at 4 °C. Following rinsing the tissue four times, the tissue was incubated in secondary antibody solution consisting of 1:1000 goat anti-rabbit biotinylated IgG (Vector Laboratories, Burlington, ON, Canada) in 0.1 M PBS for 24 hours at 4 °C. The tissue was then incubated in AB solution (Vector Laboratories) for 1 hour at room temperature. Tissue slices were then visualized with diaminobenzidine (DAB; Sigma) solution and mounted onto microscope slides, followed by dehydrated, cleared with xylene and cover-slipped with Permount (Fisher Scientific; Ottawa, ON, Canada).

#### 7.3. BrdU/NeuN double fluorescent labelling

Brain tissue was prewashed three times with 0.1 M PBS and left overnight at 4 °C. The tissue was incubated in a primary antibody solution containing 1:250 mouse anti-NeuN (Millipore; MA, USA), 0.3% Triton-X, and 3% normal donkey serum (NDS; Vector Laboratories) in 0.1 M PBS for 24 hours at 4 °C. Tissue was incubated in a secondary antibody solution containing 1:200 donkey anti-mouse ALEXA 488 (Invitrogen, Burlington, ON, Canada) in 0.1 M PBS, for 18 hours at 4 °C. After rinsed three times with PBS, tissue was washed with 4% formaldehyde, and rinsed twice in 0.9% NaCl, followed by incubation in 2N HCl for 30 minutes at 37 °C. Tissue was then incubated in a BrdU primary antibody solution consisting of 1:500 rat anti-BrdU (AbD Serotec; Raleigh, NC, USA), 3% NDS, and 0.3% Triton-X in 0.1 M PBS for 24 hours at 4 °C. Tissue was then incubated in a secondary antibody solution containing 1:500 donkey anti-rat Cy3 (Jackson ImmunoResearch; PA, USA) in 0.1 M PBS for 24 hours at 4 °C. Following three rinsed with PBS, tissue was mounted onto microscope slides and cover-slipped with PVA DABCO.

### 8. Cell counting

All counting was conducted by an experimenter blind to the group assignment of each animal using a Nikon E600 microscope. Locations of immunoreactive cells were examined whether in the dorsal or ventral dentate gyrus using the criterion defined by Banasr and others (2006), with sections 6.20-3.70mm from the interaural line defined as dorsal and sections 3.70-2.28mm from the interaural line as ventral. Cells were counted separately in each region because the dorsal hippocampus is associated with spatial learning and memory, while the ventral hippocampus is associated with emotional responses (Moser et al., 1993; Kjelstrup et al., 2002).

BrdU-immunoreactive (ir) cells were counted under a 100x oil immersion objective lens. For BrdU-ir cells, every tenth section of the hilus or the granule cell layer (GCL) that includes the subgranular zone and total immunoreactive cells per region was estimated by multiplying the aggregate number of cells per region by ten. Density of BrdU-ir cells was calculated by dividing the sum of immunoreactive cells in the hilus or GCL by volume of the corresponding region. Volume estimates of the dentate gyrus were calculated using Cavalieri’s principle by multiplying the summed areas of the dentate gyrus by distance between sections (300μm). Area measurements for the dentate gyrus were obtained using digitized images on the software ImageJ (NIH).

The percentages of BrdU/NeuN-ir cells were obtained by randomly selecting 50 BrdU-ir cells and calculating the percentage of cells that co-ir with NeuN under 40x objective lens using a Nikon E600 epifluorescent microscope (Figure. XX).

Optical density of zif268 expression in the dentate gyrus, CA3 and CA1 subregions was analyzed as an estimate of the proportion of immunoreactive cells in the subregions.

Images of the hippocampus were acquired at 100× magnification from three sections from the dorsal hippocampus and three sections from the ventral hippocampus on a Nikon E600 light microscope (Figure. 2D-I). The proportion of area that exhibited above-threshold zif268 immunoreactive intensity in the corresponding subregions was obtained using ImageJ with digitized images. The threshold was set to 2.5 times above the background gray levels. The background gray levels were the mean gray values that were obtained from five randomly selected areas without immunoactivity. The total value of optical density for each brain was calculated by dividing the total immunoreactive areas by the total area of the corresponding subregions on the three sections.

**Supplemental Fig. 1.**
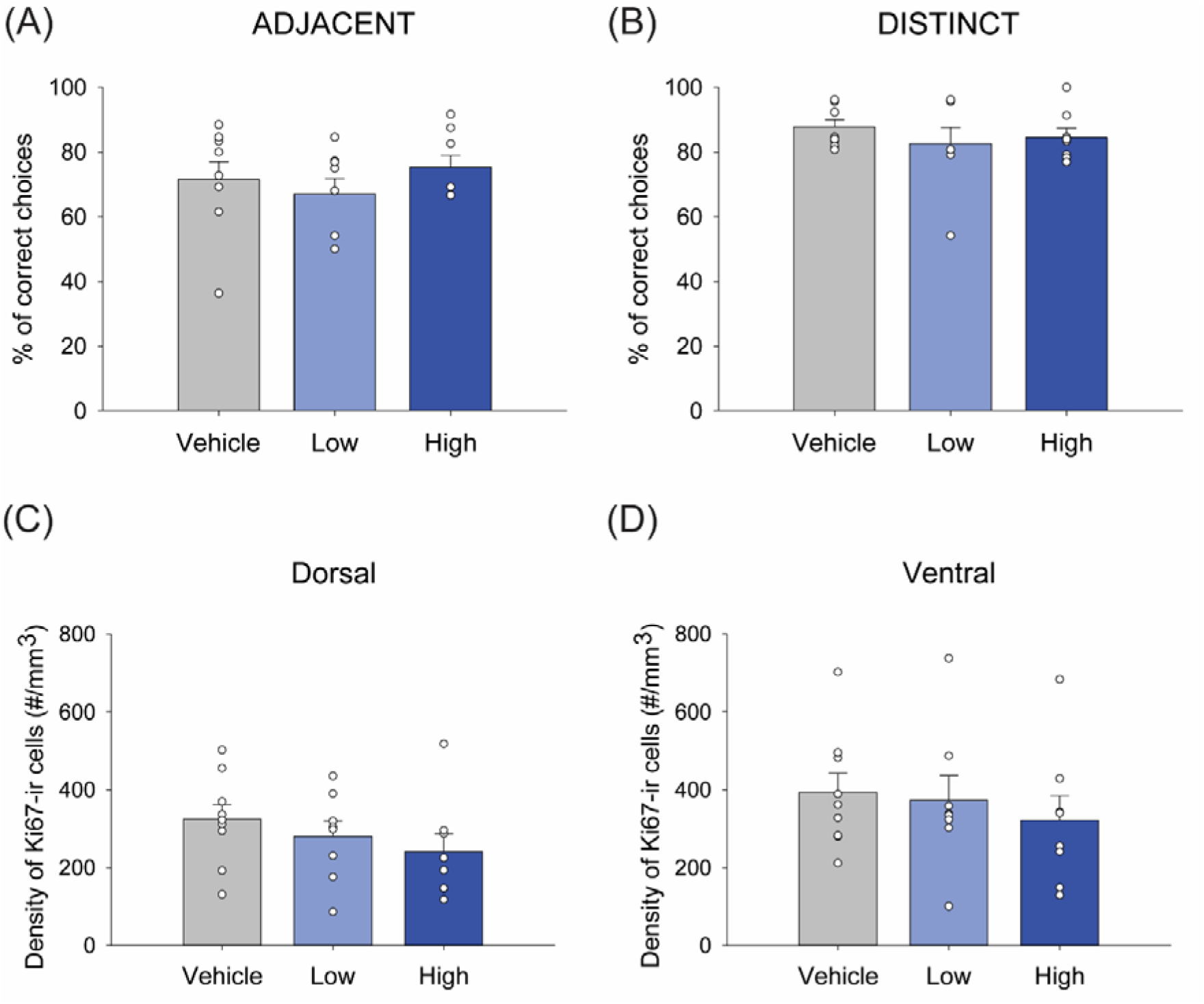
(A-B) Mean percentage of correct arm choices in ADJACENT trials (A) and DISTINCT trials (B). There were no significant main or interaction effect of treatment on the ability for spatial pattern separation. (C-D) Mean density of Ki67-immunoreactive (ir) cells in the dorsal (C) and ventral (D) dentate gyrus. There were no significant main or interaction effect of treatment on cell proliferation in the dentate gyrus. Error bars represent ± SEM. * indicates p < 0.05.

## References

Barker, J.M., Galea, L.A.M., 2008. Repeated estradiol administration alters different aspects of neurogenesis and cell death in the hippocampus of female, but not male, rats. Neuroscience 152, 888–902. https://doi.org/10.1016/j.neuroscience.2007.10.071

Becker, J.B., Rudick, C.N., 1999. Rapid effects of estrogen or progesterone on the amphetamine-induced increase in striatal dopamine are enhanced by estrogen priming: A microdialysis study. Pharmacol. Biochem. Behav. 64, 53–57. https://doi.org/10.1016/S0091-3057(99)00091-X

Chan, M., Chow, C., Hamson, D.K., Lieblich, S.E., Galea, L. a M., 2014. Effects of chronic oestradiol, progesterone and medroxyprogesterone acetate on hippocampal neurogenesis and adrenal mass in adult female rats. J. Neuroendocrinol. 26, 386–99. https://doi.org/10.1111/jne.12159

Clelland, C.D., Choi, M., Romberg, C., Clemenson, G.D., Fragniere, A., Tyers, P., Jessberger, S., Saksida, L.M., Barker, R.A., Gage, F.H., Bussey, T.J., 2009. A functional role for adult hippocampal neurogenesis in spatial pattern separation. Science (80-.). 325, 210–213. https://doi.org/10.1126/science.1173215

Dalton, M.A., McCormick, C., De Luca, F., Clark, I.A., Maguire, E.A., 2019. Functional connectivity along the anterior–posterior axis of hippocampal subfields in the ageing human brain. Hippocampus 29, 1049–1062. https://doi.org/10.1002/hipo.23097

Duarte-Guterman, P., Yagi, S., Chow, C., Galea, L.A.M., 2015. Hippocampal learning, memory, and neurogenesis: effects of sex and estrogens across the lifespan in adults. Horm. Behav. 74, 37–52. https://doi.org/10.1016/j.yhbeh.2015.05.024

Galea, L.A., Kavaliers, M., Ossenkopp, K.P., Hampson, E., 1995. Gonadal hormone levels and spatial learning performance in the Morris water maze in male and female meadow voles, Microtus pennsylvanicus. Horm. Behav. https://doi.org/10.1006/hbeh.1995.1008

Holmes, M.M., Wide, J.K., Galea, L.A.M., 2002. Low levels of estradiol facilitate, whereas high levels of estradiol impair, working memory performance on the radial arm maze. Behav. Neurosci. 116, 928–934. https://doi.org/10.1037//0735-7044.116.5.928

Kjelstrup, K.G., Tuvnes, F.A., Steffenach, H.A., Murison, R., Moser, E.I., Moser, M.-B., 2002. Reduced fear expression after lesions of the ventral hippocampus. Proc. Natl. Acad. Sci. U. S. A. 99, 10825–30. https://doi.org/10.1073/pnas.152112399

Korol, D.L., Malin, E.L., Borden, K.A., Busby, R.A., Couper-Leo, J., 2004. Shifts in preferred learning strategy across the estrous cycle in female rats. Horm. Behav. 45, 330–8. https://doi.org/10.1016/j.yhbeh.2004.01.005

Liang, X., Hsu, L.M., Lu, H., Ash, J.A., Rapp, P.R., Yang, Y., 2020. Functional Connectivity of Hippocampal CA3 Predicts Neurocognitive Aging via CA1-Frontal Circuit. Cereb. Cortex 30, 4297–4305. https://doi.org/10.1093/cercor/bhaa008

Liu, Z.Q., Vázquez-Rodríguez, B., Spreng, R.N., Bernhardt, B.C., Betzel, R.F., Misic, B., 2022. Time-resolved structure-function coupling in brain networks. Commun. Biol. 5, 1–10. https://doi.org/10.1038/s42003-022-03466-x

Marr, D., 1971. Simple Memory: A Theory for Archicortex. Philos. Trans. R. Soc. London. Ser. B, Biol. Sci. 262, 24–80.

McClure, R.E.S., Barha, C.K., Galea, L.A.M., 2013. 17β-Estradiol, but not estrone, increases the survival and activation of new neurons in the hippocampus in response to spatial memory in adult female rats. Horm. Behav. 63, 144–57. https://doi.org/10.1016/j.yhbeh.2012.09.011

Moser, E., Moser, M.B., Andersen, P., 1993. Spatial learning impairment parallels the magnitude of dorsal hippocampal lesions, but is hardly present following ventral lesions. J. Neurosci. 13, 3916–3925.

Ormerod, B. K., Falconer, E.M., Galea, L.A.M., 2003a. N-methyl-D-aspartate receptor activity and estradiol: Separate regulation of cell proliferation in the dentate gyrus of adult female meadow vole. J. Endocrinol. 179, 155–163. https://doi.org/10.1677/joe.0.1790155

Ormerod, B. K., Lee, T.T.-Y., Galea, L.A.M., 2003b. Estradiol initially enhances but subsequently suppresses (via adrenal steroids) granule cell proliferation in the dentate gyrus of adult female rats. J. Neurobiol. 55, 247–60. https://doi.org/10.1002/neu.10181

Warren, S.G., Juraska, J.M., 1997. Spatial and nonspatial learning across the rat estrous cycle. Behav. Neurosci. 111, 259–266. https://doi.org/10.1037/0735-7044.111.2.259

Wong, M., Moss, R.L., 1992. Long-term and short-term electrophysiological effects of estrogen on the synaptic properties of hippocampal CA1 neurons. J. Neurosci. 12, 3217–3225. https://doi.org/10.1523/jneurosci.12-08-03217.1992

Woolley, S., McEwen, B.S., 1994. Estradiol Regulates Hippocampal Dendritic Spine Density via an Mechanism. J. Neurosci. 74, 7680–7687.

Yagi, S., Galea, L.A.M., 2019. Sex differences in hippocampal cognition and neurogenesis. Neuropsychopharmacology 44. https://doi.org/10.1038/s41386-018-0208-4\

